# The direction of associations: prior knowledge promotes hippocampal separation, but cortical assimilation

**DOI:** 10.1101/851204

**Authors:** Oded Bein, Niv Reggev, Anat Maril

## Abstract

What does it mean to say, “a new association is learned”? And how is this learning different when adding new information to already-existing knowledge? Here, participants associated pairs of faces while undergoing fMRI, under two different conditions: a famous, highly-familiar face with a novel face or two novel faces. We examined multivoxel activity patterns corresponding to individual faces before and after learning. In the hippocampus, paired novel faces became more similar to one another through learning. In striking contrast, members of famous-novel pairs became distinct. In the cortex, prior knowledge led to integration, but in a specific direction: the representation of the novel face became similar to that of the famous face before learning, but less so vice versa, suggesting assimilation of new into old memories. We propose that hippocampal separation might resolve interference between existing and newly learned information, allowing cortical assimilation. Associations are formed through divergent but specific neural codes, that are adaptively shaped by the internal state of the system – its prior knowledge.

## Introduction

When Madonna releases a new album or starts dating a new person, one cannot avoid the deluge of media posts and ads, and so new information is added to our knowledge about Madonna. How are such new associations formed in our minds? And how is learning different when the person is less familiar to us? To be adaptive, learning cannot start de novo each time we form a new association. Rather, we cast our already existing knowledge to facilitate new learning (Alba & Hasher, 1983; Bein et al., 2015; Bransford & Johnson, 1972; Craik & Tulving, 1975; DeWitt, Knight, Hicks, & Ball, 2012; Kole & Healy, 2007; Poppenk, Kohler, & Moscovitch, 2010; Reder, Paynter, Diana, Ngiam, & Dickison, 2007; Reder et al., 2013). Indeed, robust behavioral findings demonstrate that prior knowledge facilitates memory of novel associations (e.g., Bein et al., 2015; Craik & Tulving, 1975; DeWitt et al., 2012; Fisher & Craik, 1980; Reder et al., 2013; van Kesteren, Rijpkema, et al., 2010). For example, it is easier to learn that a highly familiar person, such as Madonna, has made a new friend than to learn that two unfamiliar people have become friends (Alba & Hasher, 1983; Bein, Trzewik, & Maril, 2019; Kole & Healy, 2007; Van Overschelde & Healy, 2001). However, while existing memory representations can serve as scaffolding for the assimilation of new information (Ghosh & Gilboa, 2014; Gilboa & Marlatte, 2017; McClelland, 2013; Tse et al., 2007, 2011; van Kesteren, Rijpkema, Ruiter, & Fernandez, 2010), they can also at the same time produce interference, as multiple existing associations may render the acquisition of new memories more difficult (Anderson, 1974, 1983; Reder & Anderson, 1999; Reder et al., 2007). In the face of this conundrum, the question arises: when forming a new memory, how do we successfully utilize prior knowledge while also protecting against interference?

One possibility is a division of labor, such that prior knowledge biases the *cortical* memory system towards assimilation of novel information (Gilboa & Marlatte, 2017), while shifting *hippocampal* processes to resolve interference (McClelland, McNaughton, & Oreilly, 1995). Across rodents and humans, studies demonstrate that during new learning, prior knowledge enhances cortical activation and cortico-cortical functional connectivity, while also modulating hippocampal activation and hippocampal functional connectivity with the cortex (Amer, Giovanello, Nichol, Hasher, & Grady, 2019; Bein, Reggev, & Maril, 2014; Brod, Lindenberger, Wagner, & Shing, 2016; Kumaran, Summerfield, Hassabis, & Maguire, 2009; Liu, Grady, & Moscovitch, 2016; Reggev, Bein, & Maril, 2016; Sommer, 2017; Tse et al., 2007, 2011; van Buuren et al., 2014; van Kesteren, Fernandez, Norris, & Hermans, 2010; van Kesteren, Rijpkema, et al., 2010; Wagner et al., 2015; Wang, Tse, & Morris, 2012; for reviews, see Gilboa & Marlatte, 2017; Preston & Eichenbaum, 2013; van Kesteren, Ruiter, Fernández, & Henson, 2012). Critically, however, univariate activation and functional connectivity studies cannot address the content of learning, so the question of *how* prior knowledge modulates the neural representation of novel associations in these memory systems remains open. As such, in the present study, we asked whether the cortical system might support the beneficial effects of prior knowledge on novel learning through assimilation, while the hippocampus defends against potential interference from the same knowledge.

Theoretical frameworks propose that hippocampal processes mitigate interference between novel memories and existing memories (McClelland et al., 1995). Indeed, univariate activation in the hippocampus during learning of a novel association reduces forgetting of a previously-learned, related association (Kuhl, Shah, DuBrow, & Wagner, 2010). But how does the hippocampus resolve interference between competing memories? Research in humans and rodents has shown that the hippocampus separates overlapping memories by allocating a distinct activity pattern to each in a process known as pattern separation (Baker et al., 2016; Bakker, Kirwan, Miller, & Stark, 2008; Chanales, Oza, Favila, & Kuhl, 2017; Favila, Chanales, & Kuhl, 2016; LaRocque et al., 2013; Leutgeb, Leutgeb, Moser, & Moser, 2007). This process could potentially mitigate proactive interference from existing associations when a novel association is added to a previously known item. If so, we would expect that learning about Madonna’s new friend would cause the representations of Madonna and her friend to become more distinct^1^.

While prior knowledge might bias the hippocampus towards pattern separation, it might also lead to assimilation of novel information in cortical regions (McClelland, 2013; van Kesteren et al., 2012). As noted above, prior knowledge increases cortical activation and cortico-cortical functional connectivity (Liu et al., 2016; Maril et al., 2011; Reggev et al., 2016; Staresina, Gray, & Davachi, 2009; van Kesteren, Rijpkema, et al., 2010). While this supports cortical involvement in prior knowledge effects on learning, it remains unclear precisely how assimilation occurs at the level of neural representation. If new information is indeed woven into an existing cortical representation (Ghosh & Gilboa, 2014; Gilboa & Marlatte, 2017), we propose that when supported by prior knowledge, the learning of novel associations leads to asymmetric cortical effects. To illustrate, since a cortical representation of Madonna has been stabilized over a lifetime of exposure, it might only change slightly to incorporate the novel association with her friend. The representation of the friend, however, will undergo a disproportionately larger transformation during learning, in order to be assimilated into our stronger, existing representation of Madonna.

To test these ideas, we had human participants learn associations between different pairs of faces. One type of pair comprised a famous face and a novel face, whereas the other type of pair comprised two novel faces. Critically, we presented each face alone both before and after associative learning (Kim, Norman, & Turk-Browne, 2017; Schapiro, Kustner, & Turk-Browne, 2012; Schlichting, Mumford, & Preston, 2015). Using a representational similarity analysis (RSA; Kriegeskorte, Goebel, & Bandettini, 2006; Kriegeskorte, Mur, & Bandettini, 2008), we assessed how the multivoxel-pattern representations of individual faces changed as a function of learning and of whether or not they included an association with a famous face; that is, whether or not the pair contained an element of prior knowledge. A final associative memory test was used to determine whether learning-related representational changes contributed to subsequent memory. We show that prior knowledge differentially modulates associative learning in the hippocampus versus the cortex. Namely, prior knowledge leads to representational separation in the hippocampus, but assimilation in the cortex. Together, these findings suggest a candidate mechanism for assimilating new information into existing knowledge structures while reducing memory interference.

## Results

During the associative learning task, participants repeatedly observed two types of face pairs: either a famous face and a novel face (Prior Knowledge, PK) or two novel faces (No Prior Knowledge, n-PK). Before and after learning, each face was presented alone on the screen to enable capture of its multivoxel pattern representation (Methods, Figure 1). In both tasks, participants performed orthogonal gender judgments about the faces. Accuracy during learning and during the pre- and post-learning scans was above 96%, demonstrating that participants complied with task instructions (see Supplementary Information for further details).

**Figure 1.**
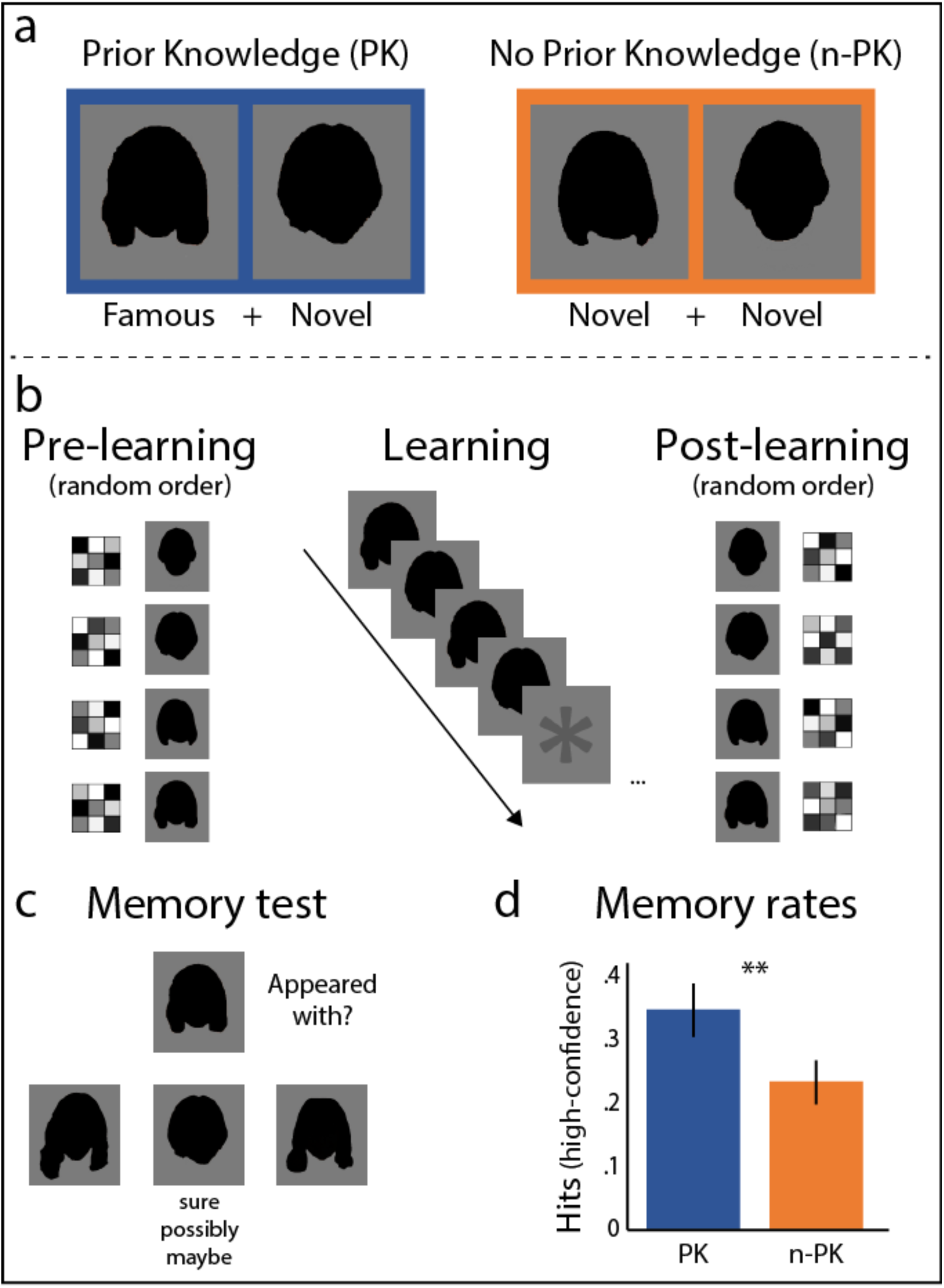
Design and behavior. a) Experimental conditions: participants learned pairs of faces, either a famous and a novel face in the Prior Knowledge condition (PK), or two novel faces in the No Prior Knowledge condition (n-PK). Note: faces are blackened in this figure for publication, to avoid inclusion of identifying information about people. Participants in the experiment viewed actual faces. b) While in the scanner, participants viewed the pairs 12 times in 12 cycles; each cycle included all pairs in a random order. Before and after associative learning, participants in the scanner also viewed each face presented alone, in random order. This allowed us to capture the multivoxel activity pattern of each face, for pattern similarity analysis (see Methods). c) After the post-similarity scan, participants performed an associative memory test, in which they indicated which of the three bottom faces appeared with the face on the top and rated their confidence (sure/possibly/maybe). d) Behavioral

### Associative memory test

After the post-learning scan, participants were tested for associative memory for all face pairs in a three-alternative forced choice task. Accuracy rates for both types of face pairs (PK, n-PK) were significantly above chance (33%; PK: *M* = .46, *SD* = .18, *t_(18)_* = 2.89, *p* = .01, Cohen’s *d* = .66; n-PK: *M* = .44 *SD* = .14, *t*_(18)_ = 3.24, *p* = .005; Cohen’s *d* = .74). Overall accuracy rates did not reliably differ between pair types (*t*_(18)_ = .41, *p* = .69). Importantly, participants had more high-confident hits (“sure” and “possibly” responses, excluding “maybe” responses) for PK pairs compared to n-PK pairs (PK: *M* = .35, *SD* = .18, n-PK: *M* = .23, *SD* = .15, *t*_(18)_ = 2.88, *p* = .01, *Cohen’s d* = .66; Figure 1d). Thus, our results are consistent with previous findings showing that prior knowledge enhances new learning (Bein et al., 2015, 2019; Kole & Healy, 2007; Liu et al., 2016; Van Overschelde & Healy, 2001).

### Hippocampus: Prior knowledge leads to representational separation

We tested whether the hippocampal representations of two associated items become more distinct when learning involved prior knowledge. Due to the known role of the hippocampus in supporting associative memory (e.g., Davachi, 2006; Diana, Yonelinas, & Ranganath, 2007; Eichenbaum, 2004; Squire, 2004), we predicted that representational changes would be specific to remembered face pairs. To that end, we examined how learning altered the similarity between the multivoxel BOLD activity patterns of items in a pair by computing the change in similarity from before to after learning. We then compared learning-related changes in similarity between members of PK and n-PK pairs, and dependent on whether the association between faces was later remembered (high-confidence hits) or forgotten in the final associative memory test (Figure 2, left).

**Figure 2.**
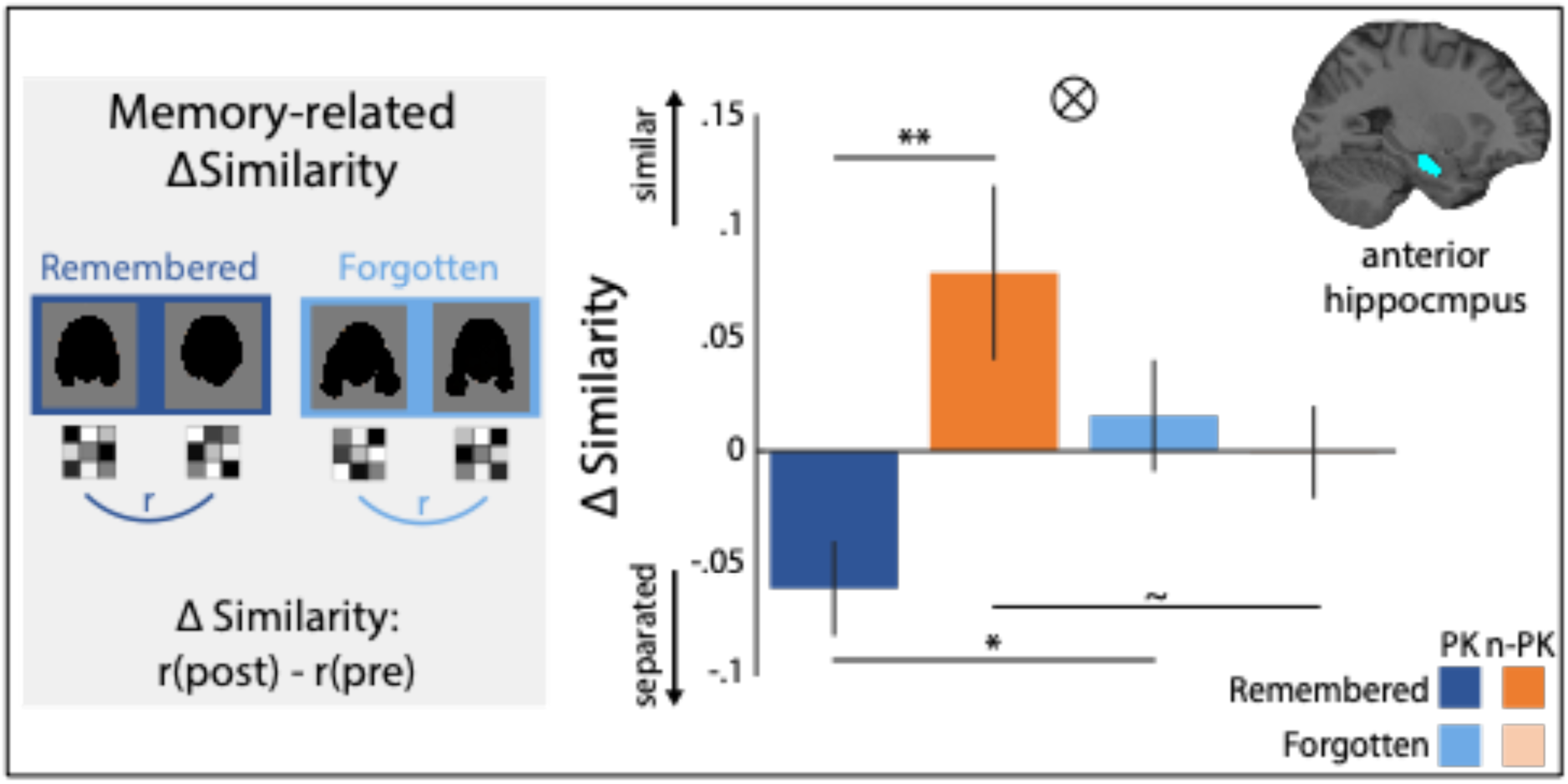
Prior knowledge and memory-dependent representational changes in the hippocampus. Left: Pairs were grouped based on Memory in the later associative memory test (remembered pairs include high-confidence hits), and Prior Knowledge (PK/no prior knowledge, n-PK). Note: faces are blackened in this figure for publication, to avoid inclusion of identifying information about people. Participants in the experiment viewed actual faces. The multivoxel activity patterns of items in each pair were correlated before and after learning, and the pre-to post-learning difference in correlation values (Fisher-transformed) was calculated. Right: Results from the left anterior hippocampus. Similarity after learning was lower between members of PK pairs, in contrast to increase in similarity in n-PK pairs. Similarity differences were specific to remembered pairs. \*\**p* < .01, \**p* < .05, ∼*p* < .1. Interaction: *p* < .006

Similarity differences were submitted to a 2 (Prior Knowledge: PK, n-PK) by 2 (Memory: remembered – high-confidence hits only, forgotten) repeated-measures ANOVA. As shown in Figure 2, there was a significant interaction between Prior Knowledge and Memory in the left anterior hippocampus (*F*_(1,17)_ = 10.12, *p* = .0055, η*_p_*^2^ = .37^2^). This interaction stemmed from face representations in PK pairs becoming more *distinct* (less similar) from one another after learning, in contrast to representations in n-PK pairs, which became *more* similar to each other after learning (remembered, PK vs. n-PK: *t*_(17)_ = 2.98, *p* = .008, *Cohen’s d* = .70). Interestingly, similarity changes only occurred for pairs that were later remembered (significant for PK pairs, remembered vs. forgotten: *t*_(17)_ = 2.48, *p* = .02, *Cohen’s d* = .58; approaching significance for n-PK pairs, remembered vs. forgotten: *t*_(17)_ = 1.92, *p* = .07, *Cohen’s d* = .45).

For robustness, and to further ensure that these effects were specific to pairs that were learned together, rather than a general effect of observing faces with prior knowledge versus without prior knowledge, we compared remembered pairs to shuffled pairs (items from the same PK/n-PK pair-type that did not appear together at learning, see Methods). Once again, remembered PK pairs were significantly less similar to each other than were PK-shuffled pairs (*t*_(17)_ = 3.05, *p* = .007, *Cohen’s d* = .72; PK-shuffled: *M* = -.004, *SD* = .04). Remembered n-PK pairs were more similar than n-PK shuffled pairs (n-PK: *t*_(17)_ = 2.09, *p* = .052, *Cohen’s d* = .49; n-PK-shuffled: *M* = -.001, *SD* = .015). Thus, in the n-PK condition, our results are consistent with previous research demonstrating increased similarity for novel pairs of items (Schapiro et al., 2012). Critically, our results show the opposite pattern when novel information becomes associated with prior knowledge due to learning – in this case, the items’ representations became more separated.

### Prior knowledge enhances hippocampal-cortical functional connectivity

If semantic knowledge is represented in the cortex (Binder & Desai, 2011; Gilboa & Marlatte, 2017; Gobbini & Haxby, 2007) and hippocampal processes resolve interference between this knowledge and new learning, we reasoned that there should be higher hippocampal-cortical functional connectivity during learning in PK pairs compared to n-PK pairs. Such crosstalk might reflect input of cortical information about the famous faces to the hippocampus, or top-down control signaling the need for interference resolution. To test this, we examined functional connectivity between the left anterior hippocampus, where multivoxel-pattern similarity differences were observed, and the rest of the brain using a psychophysiological interaction analysis (PPI; McLaren, Ries, Xu, & Johnson, 2012). We compared all PK pairs to n-PK pairs during the associative learning task. Consistent with our prediction, we found a host of cortical regions that demonstrated higher functional connectivity with the anterior hippocampus for PK compared to n-PK pairs, including the left inferior frontal gyrus, the angular gyrus, and the left middle temporal gyrus (see Figure 3 and Supplementary Table 1 for detailed results)^3^. All of these regions are involved in the processing of famous faces, or more broadly in semantic processing (Binder & Desai, 2011; Binder, Desai, Graves, & Conant, 2009; Gobbini & Haxby, 2007; Natu & O’Toole, 2011). No significant differences in functional connectivity were observed in the opposite n-PK > PK contrast. These results show that prior knowledge enhances communication between cortical regions and the hippocampus during new associative learning, consistent with our predictions.

**Figure 3.**
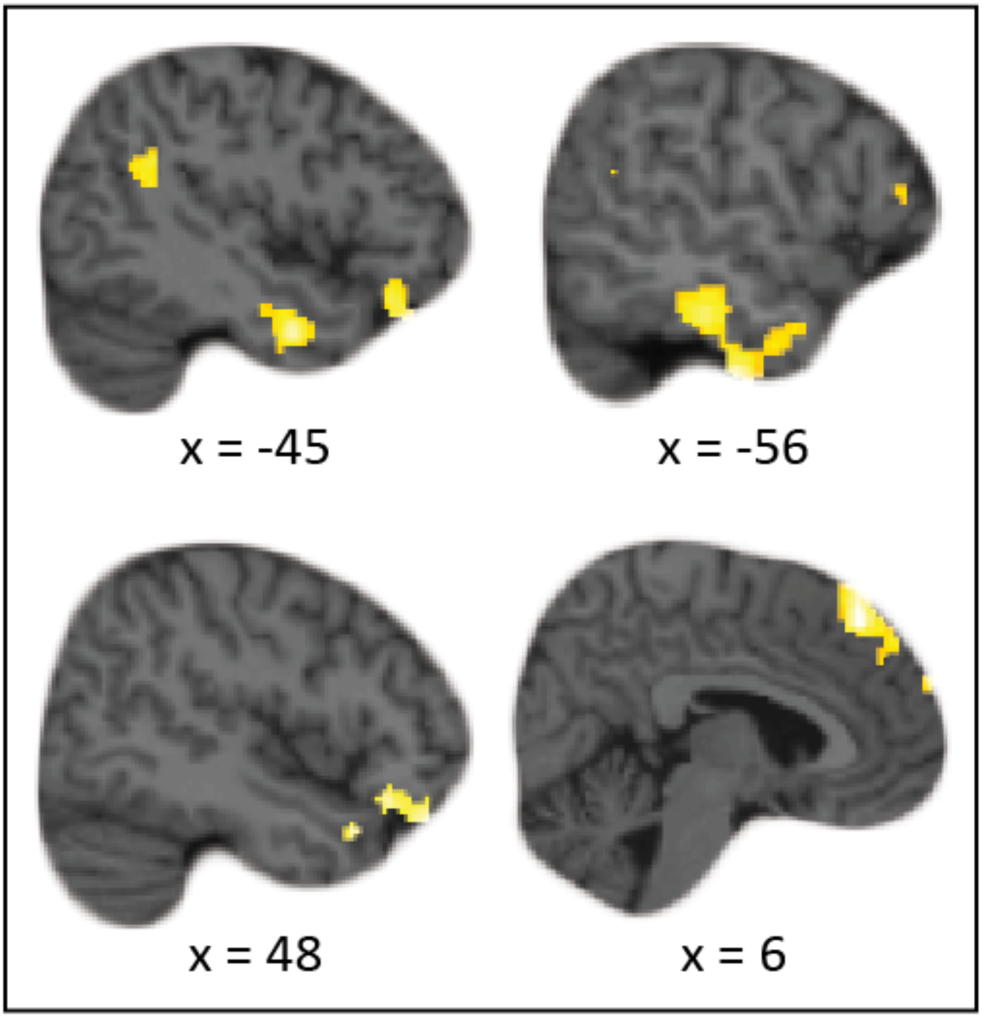
Regions demonstrating significantly higher functional connectivity (PPI) with the left anterior hippocampus for prior knowledge (PK) pairs compared to no prior knowledge (n-PK) pairs during associative learning (see also Table S1).

### Cortex: New information is assimilated into cortical knowledge structures

Next, we tested whether prior knowledge led to assimilation in the cortex (McClelland, 2013; Sharon, Moscovitch, & Gilboa, 2011; Tse et al., 2007, 2011; van Kesteren et al., 2012). Specifically, we asked whether cortical representations showed evidence of asymmetric updating, in which the representation of a novel face after learning became more similar to the original (pre-learning) representation of the famous face it was associated with. This could indicate that the representation of the novel face was woven into the representation of the famous face (see Introduction, Methods). To the extent that assimilation co-occurs with, or even depends on, hippocampal interference-resolution, we reasoned that this should be observed in regions that communicated with the hippocampus during learning. Thus, we chose an ROI that showed higher functional connectivity with the anterior hippocampus for PK pairs than for n-PK pairs (functional connectivity analysis, Figure 3). We focused on the left inferior frontal gyrus (left IFG) due to its roles in mediating the effects of prior knowledge on new learning (Maril et al., 2011; Reggev et al., 2016; Staresina et al., 2009) and in semantic processing more broadly (Badre, Poldrack, Paré-Blagoev, Insler, & Wagner, 2005; Badre & Wagner, 2007; Binder & Desai, 2011). Indeed, we found that in this region, associated items became more similar to one another from pre- to post-learning than did shuffled pairs (across PK and n-PK pairs), indicating associative learning (see Supplementary Material).^4^

When comparing the representational similarity of items after learning, it is unclear whether similarity increases reflect changes in the representations of both items, such that they both become more similar to each other (symmetric changes), or in one item, which becomes more similar to the other (asymmetric changes). To test our main hypothesis regarding asymmetry in learning, we reasoned that comparing the post-learning pattern of the novel B-face to the pre-learning pattern of its paired famous A-face would give us a pure measure of learning-related changes in the novel B-face (we performed an identical procedure for novel-novel n-PK pairs). Thus, we compared the representational similarity between the B-face after learning (always novel) and the A-face (famous or novel) before learning, as well as the representational similarity between the A-face after learning and that of the B-face before learning. We then subtracted the latter from the former to get an asymmetry measure: if both face representations became equally more similar to one another, there would be no difference between these two values. If, however, the B-face after learning became more similar to the A-face before learning, while the A-face did not become more similar to the B-face before learning, then we should observe a positive value, indicating asymmetry (Schapiro et al., 2012; Methods and Figure 4). Consistent with our prediction, positive and significant asymmetry was observed only for PK pairs (compared to zero: *t*_(18)_ = 2.71, *p* = .01, *Cohen’s d* = .62; or compared to asymmetry in the shuffled pairs: *t*_(18)_ = 3.53, *p* = .002, *Cohen’s d* = .81; Figure 4). No asymmetry was found for n-PK pairs (either relative to zero or to asymmetry in shuffled pairs, *t*’s < .25, *p*’s > .8).

**Figure 4.**
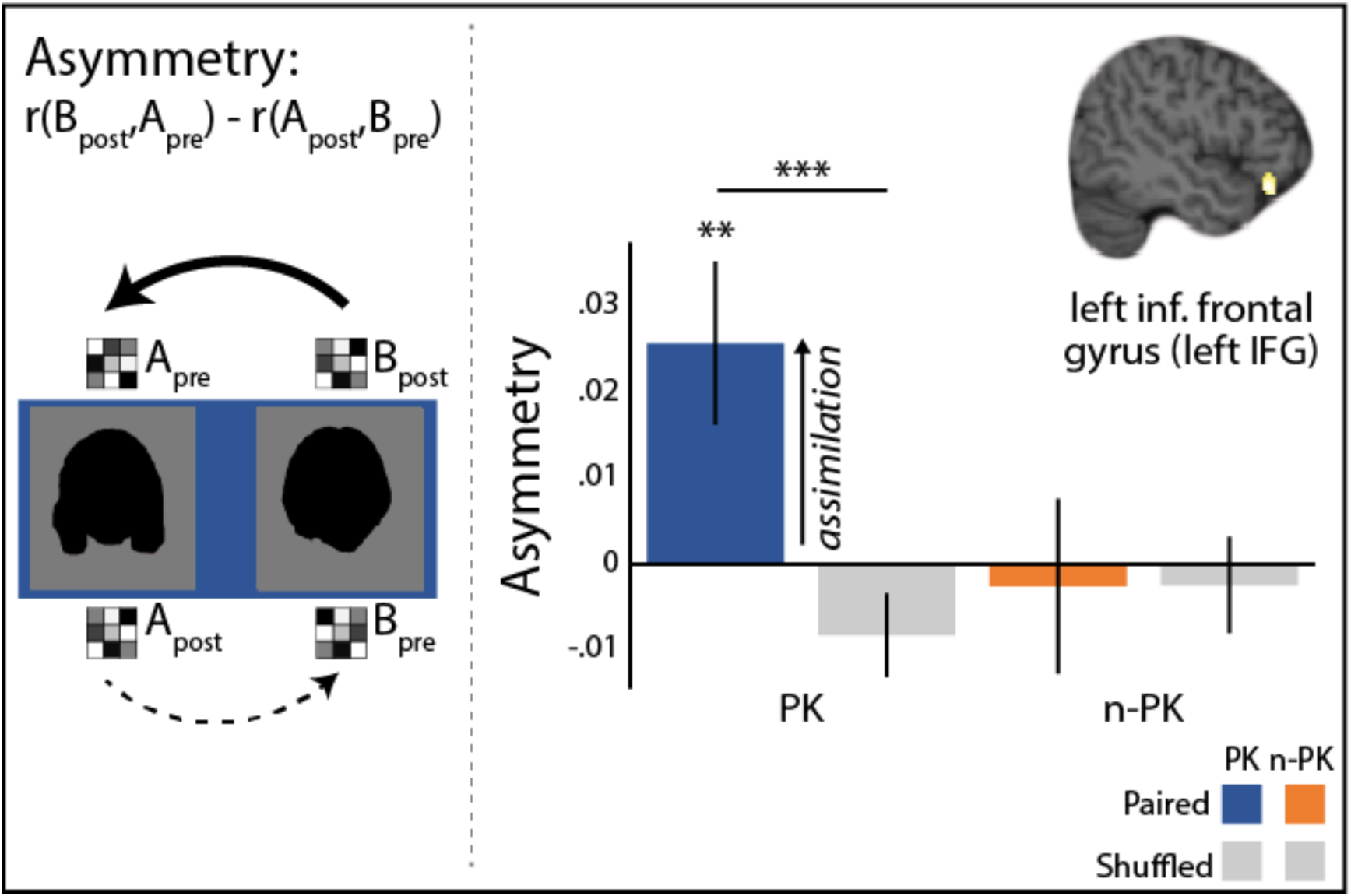
Similarity changes in the left inferior frontal gyrus (left IFG). Left: asymmetry in learning reflects the extent to which the representation of the B-face after learning became more similar to that of the A-face before learning than the representation of the A-face became similar to that of the B-face (B_pre_). In accordance, the multivoxel activity pattern of the B-face post-learning (B_post_) was correlated with the pattern of the A-face pre-learning (A_pre_), and the pattern of the A-face post-learning (A_post_) was correlated with the pattern of the B-face pre-learning (B_pre_). Then, the latter similarity value was subtracted from the former as shown above, to reflect the extent to which the B-face became more similar to the A-face than the A-face became to the B-face. We interpreted asymmetry as assimilation of the B-face into the representation of the A-face. Note: faces are blackened in this figure for publication, to avoid inclusion of identifying information about people. Participants in the experiment viewed actual faces. Right: asymmetry was observed in the left inferior frontal gyrus in the prior knowledge (PK) condition for paired faces only, but not in the no prior knowledge (n-PK) condition. \*\*\**p* < .005, \*\**p* = .01

### Ruling out alternative explanations

We emphasize that our results are unlikely to reflect some general response to famous faces during the pre-or post-learning scans. First, we report the difference between the pre and post scans, and pattern similarity was computed with famous faces in both. Second, critically, our control comparisons in all of the similarity analyses were within condition: shuffled pairs in the PK condition only included pairs in the PK condition, and likewise for the n-PK condition. Nevertheless, the results were specific to paired faces. Another possibility is that the novel faces that were paired with famous faces in PK pairs carry with them some unique status due to becoming associated with famous faces. However, again, any such status that is unrelated to associative learning would have been observed in the shuffled pairs as well. Regarding the hippocampal results, pairs were separated to remembered versus forgotten associations within each pair type, and our results were specific to the remembered faces, alleviating the above concerns for the memory analysis as well.

We further controlled for potential differences in univariate activation during the pre- and post-learning scans (Coutanche, 2013; Davis & Poldrack, 2013; LaRocque et al., 2013). As mentioned above, differences in univariate responses between PK and n-PK pairs, if they arose, should have influenced paired and shuffled items alike. Nevertheless, we performed an additional control analysis, in which we included the univariate activity together with the factors of Prior Knowledge (PK/n-PK) and Pairing (and Memory, where relevant) in a multiple linear regression. All of the analyses reported here hold when controlling for univariate activation (see Supplementary Material). Thus, our representational similarity effects are unlikely explained by univariate activity.

## Discussion

Here, we asked how new associations are represented in the human brain, and how are individual associations different when adding new to old memories versus learning thoroughly novel associations. Decades of behavioral research have shown the power of prior knowledge to facilitate new learning, such that adding a novel piece of information to existing knowledge is typically easier than thoroughly new learning (Alba & Hasher, 1983; Bein et al., 2015, 2019; Bransford & Johnson, 1972; Craik & Tulving, 1975; DeWitt et al., 2012; Kole & Healy, 2007; Long & Prat, 2002). In recent years, research has shown that prior knowledge increases cortical activation and functional connectivity, and modulates hippocampal activation and hippocampal-cortical functional connectivity (for reviews, see Ghosh & Gilboa, 2014; Gilboa & Marlatte, 2017; Preston & Eichenbaum, 2013; van Kesteren et al., 2012). Critically, as previous studies only looked at the strength of brain activation and lesions, they could not address the content of learning – namely, what learning processes take place?

Theoretically, prior knowledge could facilitate new learning through multiple processes. Prior knowledge is thought to serve as scaffolding for learning by providing an existing cortical representation into which new information can be assimilated. However, existing associations can also interfere with new learning, causing difficulty to associate an item with novel information (i.e., “fan effect”, Anderson, 1974, 1983; Reder & Anderson, 1999; Reder et al., 2007). Thus, prior knowledge may promote mechanisms aimed at mitigating interference, such as hippocampal pattern separation. To address this possibility, we examined how prior knowledge altered the neural representations of newly learned associations between either a famous and a novel face or two novel faces.

We found that prior knowledge led to greater separation of underlying neural representations in the hippocampus. Multivoxel activity patterns of members of famous-novel pairs became less similar to each other after associative learning, whereas representations of novel-novel face pairs became more similar to each other after associative learning. These learning-dependent changes in similarity were specific to face pairs that participants later remembered, and did not occur for forgotten pairs. In contrast, prior knowledge led to cortical assimilation, expressed in asymmetric representational changes in the left inferior frontal gyrus. Specifically, we found that the neural representations of novel faces following learning became more similar to the representations of their associated famous faces before learning. Together, these findings show a flexible and directional creation of associations in the human brain, that is specific to memory systems and is highly determined by the state of prior knowledge.

### Shifting the direction of hippocampal learning

We found that associative learning processes in the hippocampus were highly dependent on prior knowledge. Consistent with previous findings, the representations of novel pair members became more similar to one another after associative learning (Schapiro et al., 2012). In contrast, we found that famous-novel pairs became more separated after learning. This increased separation supports the idea of interference resolution, in line with previous research on pattern separation (e.g., Bakker et al., 2008; Chanales et al., 2017; Favila et al., 2016; Leutgeb et al., 2007). Importantly, separation mediated successful associative memory specifically when prior knowledge was involved and there was a need to overcome interference from previous associations. Future research can investigate precisely how previous associations interfere with new learning, and how separation processes might facilitate resolution of this interference.

The exact manner by which prior knowledge drives the hippocampus towards separation versus similarity, or integration, is currently unknown. One possibility is that top-down control signals modulate hippocampal computations during new learning. A wealth of research suggests that prefrontal-hippocampal interactions mediate cognitive control processes that select representations for encoding or retrieval from memory (see e.g., Eichenbaum, 2017; Simons & Spiers, 2003, for reviews). These control processes might promote interference resolution by modifying hippocampal representations (Benoit, Hulbert, Huddleston, & Anderson, 2015; Guise & Shapiro, 2017; Rajasethupathy et al., 2015; Shimamura, Jurica, Mangels, Gershberg, & Knight, 1995). Supporting this possibility, we saw prior knowledge lead to greater hippocampal-prefrontal interactions during learning and greater subsequent separation in hippocampal representations thereafter.

Another possibility is that differential neuromodulatory input to the anterior hippocampus (Kafkas & Montaldi, 2018; Rangel-Gomez & Meeter, 2016) biases the hippocampus towards separation versus similarity (Duncan & Schlichting, 2018; Giocomo & Hasselmo, 2007; Hasselmo, Bodelón, & Wyble, 2002). Kafkas and Montaldi (2018) recently proposed that different types of novelty, such as absolute or contextual, are both detected in the anterior hippocampus, but with different neurotransmitters mediating each type. In our study, novel-novel pairs may evoke an absolute novelty signal, because neither image has ever been seen before. In contrast, new associations involving prior knowledge might promote a contextual novelty signal, because the novel face is novel in the context of the highly-familiar face. It has been proposed that absolute novelty enhances acetylcholine input to the hippocampus, while contextual novelty involves the release of dopamine and norepinephrine (Kafkas & Montaldi, 2018; see also Hasselmo, Wyble, & Wallenstein, 1996; Lisman & Grace, 2005; Meeter, Murre, & Talamini, 2004). Different neurotransmitters might further lead to separation versus similarity in the hippocampus (e.g., Duncan & Schlichting, 2018; Giocomo & Hasselmo, 2008; Hansen & Manahan-Vaughan, 2015; Harley, 2007), as was observed here.

While we found neural signatures of both separation and integration in the anterior hippocampus, it has recently been proposed that these different computations may be localized to the posterior versus anterior hippocampus, respectively (Brunec et al., 2018; Poppenk, Evensmoen, Moscovitch, & Nadel, 2013). Following a previous recent study showing separation in the anterior hippocampus (Tompary & Davachi, 2017), we interpret our findings within a framework embedding the hippocampus in a larger functional network. Anatomically, the anterior hippocampus receives preferential input from the perirhinal cortex (via entorhinal cortex; Burwell, 2000; Suzuki & Amaral, 1994). The perirhinal cortex supports conceptual semantic knowledge and is involved in the processing of items and their features (Barense et al., 2012; Brown & Aggleton, 2001; Clarke & Tyler, 2014; Davachi, Mitchell, & Wagner, 2003; Staresina & Davachi, 2008; Staresina, Duncan, & Davachi, 2011), potentially as a part of a larger antero-temporal network (Ranganath & Ritchey, 2012). In this context, it is not surprising that in our study, which involved associating items and incorporated semantic knowledge, we found representational changes in the anterior hippocampus. Our findings thus suggest that the anterior hippocampus might mediate both separation and integration, dependent on internal knowledge.

The type of prior knowledge involved in new learning could be critical in shifting the hippocampus towards a separation mode versus an integration mode. Using a transitive-inference paradigm, a previous study showed that after an A-B pair was learned, learning of an overlapping A-C pair resulted in greater pattern similarity between the B and the C items in the anterior hippocampus (Schlichting et al., 2015, letters represent different items). Although they used merely visual associations, conceptualizing the A-B association as some prior knowledge to which a novel A-C association is added, our separation finding might seem to diverge from these previous similarity results. This might point towards factors that bias hippocampal representations. For example, the A-B pairs form a single association learned over only a few repetitions, whereas the knowledge about the famous faces that we used in the current study is highly learned and involves a rich network of strong associations. Thus, one or a few weaker prior associations as in the case of the A-B pairs might not interfere with new learning and result in similarity (Schlichting et al., 2015), while multiple strong associations require interference-resolution and necessitate separation.

Another potentially interesting factor is consolidation, or time. Here, we used knowledge that was acquired long before our study and was well consolidated, while in the transitive inference study, the A-B associations were learned immediately before the A-C associations. Indeed, a 24-hour delay between learning the old A-B and the novel A-C association reduces transitive inference (Zeithamova & Preston, 2017). Previous studies have further shown that prior knowledge established immediately before new learning does not enhance associative memory to the same extent as long-held, pre-existing knowledge (Poppenk & Norman, 2012). Future research should elucidate how the time difference between the initial acquisition of knowledge and the addition of novel associations modulates associative learning in the hippocampus.

### Asymmetric cortical learning

Cortical learning has fascinated researchers across domains, asking about the processes as well as the timeline that characterizes learning in the cortex (e.g., Ahissar & Hochstein, 2004; Binder & Desai, 2011; Brodt et al., 2018, 2016; Hebscher, Wing, Ryan, & Gilboa, 2019; Heeger, 2017; McClelland, McNaughton, & Oreilly, 1995; Moscovitch, Cabeza, Winocur, & Nadel, 2016; Tse et al., 2011). Here, we investigated the content of individual associations in the cortex (Tompary & Davachi, 2017), and whether the existence of knowledge changes the direction of association, in a specific and predictable way. Theoretical accounts suggest that through learning, new information is assimilated into cortical knowledge structures (Ghosh & Gilboa, 2014; McClelland, 2013; van Kesteren et al., 2012). While previous empirical findings of cortical activation and functional connectivity support this view (Liu et al., 2016; Maril et al., 2011; Reggev et al., 2016; Staresina et al., 2009; van Kesteren, Rijpkema, et al., 2010), they do not address the content of learning. Here, we found that in the left inferior frontal gyrus, representations of the novel faces after learning became more similar to representations of the famous faces before learning (see Methods and Figure 4), more so than representations of the famous faces after learning became to representations of the novel faces before learning. Critically, we did not observe such asymmetry in learning when both faces were novel. We propose that asymmetry in learning reflects the assimilation of new information into existing knowledge. This semantic knowledge is acquired across multiple encounters, so existing representations are not modified as strongly when learning additional novel information. Meanwhile, representations of novel information undergo large transformations as they become woven into existing schemas.

Cortical assimilation might be influenced by whether novel information is consistent, inconsistent, or arbitrary with respect to prior knowledge. Here, we show assimilation based on arbitrary associations. Thus, our results are consistent with theories implicating assimilation as a general mechanism for knowledge-supported learning (Alba & Hasher, 1983; Bellezza & Buck, 1988; Ghosh & Gilboa, 2014; Gilboa & Marlatte, 2017; Sharon et al., 2011). Note, however, that others have proposed that assimilation specifically mediates learning of information that is consistent with prior knowledge (McClelland, 2013; Tse et al., 2007; van Kesteren et al., 2012). Similar to our view, these latter frameworks rely on the assumption that the hippocampus is required to prevent interference between new information and cortical knowledge (McClelland et al., 1995). Thus, when novel information is consistent and elicits less conflict with cortical knowledge structures, cortical assimilation can occur, and hippocampal involvement is reduced (McClelland, 2013; van Kesteren et al., 2012). In the current study, we show that cortical assimilation can occur in parallel with hippocampal involvement. We thus propose that interference resolution occurs either because the novel information elicits less interference to begin with, as in the case of schema-consistent information, or because the hippocampus contributes to the resolution of interference, as in our study. How neural systems may co-operate to determine the neural representation of new memories, and how these processes are shaped by consistency with prior knowledge, are fascinating questions for future research (Preston and Eichenbaum, 2013).

While we interpret our asymmetry finding as assimilation, it is also possible that after learning, the second face in the pair (B-face) brings to mind the first face (A-face) more so than vice-versa. A previous study found that after sequential learning, the representation of the *first* item in a pair following learning became more similar to the representation of the *second* item in the pair before learning (Schapiro et al., 2012). This was interpreted as the first item bringing to mind, or predicting, the second item due to their temporal contingency. In contrast, we found that the second face after learning became similar to the first face before learning.

However, applying Schapiro et al.’s (2012) interpretation to our findings does raise the possibility that the asymmetry effect we observed reflects the B-face bringing to mind the A-face. We find this interpretation less likely in our case, because asymmetry was observed in the *opposite* direction from the temporal order of learning. Moreover, for asymmetry to arise, the B-face should elicit the A-face more so than the A-face elicits the B-face. While intriguing, it seems less reasonable that the B-face, both novel and temporally second, would make a stronger cue than the A-face, which is a famous face and was temporally first in the pair during learning. We thus find the assimilation interpretation more plausible, but acknowledge that the alternative retrieval interpretation should be tested.

### Concluding remarks

We asked what does it mean to say “an association was created”? Importantly, an adaptive learning system does not start any learning experience tabula-rasa, but rather it utilizes what it already knows about the world. However, reliance on prior knowledge is a double-edged sword, as existing memories can enhance but also impair new learning (Anderson, 1974, 1983; Alba & Hasher, 1983; Atienza, Crespo-Garcia, & Cantero, 2011; Bein et al., 2015; DeWitt et al., 2012; Kole & Healy, 2007; Long & Prat, 2002; Reder & Anderson, 1999; Reder et al., 2007). While many important questions remain open, our findings suggest a novel putative mechanism for learning as it typically occurs in our every-day lives: we usually add new information to what we already know. In this case, we propose that new information is assimilated into our prior knowledge in the cortex, while hippocampal pattern separation mitigates interference between new and old memories. Thus, the current study clearly demonstrates that associative learning is flexible and directional, specific to memory systems, and highly dependent on prior knowledge.

## Author Contributions

O.B., N.R., and A.M designed the experiment; O.B. collected and analyzed the data; A.M. provided guidance; O.B., wrote the initial draft of the paper; O.B., N.R., and A.M. revised and edited the paper; A.M. acquired funding

## Acknowledgements

This work was supported by Israel Science Foundation (150/16, to A.M.). We thank Haim Cohen and Alexa Tompary for facilitating the fMRI analyses and Maayan Trzewik for her help in data collection. We further thank Alexa Tompary, David Clewett, Tarek Amer, Catherine Hartley, and Katherine Nussbaum for their insightful comments and suggestions in preparing the manuscript.

## Methods

### Participants

Nineteen right-handed native Hebrew-speakers participated in the study (nine women; mean age: 26.94 years, range: 22-31 years). Five additional participants were excluded from the analysis: two due to excessive movement (more than 3mm across all pre-learning, post-learning, and associative learning scans); two due to insufficient knowledge about the famous faces, as defined by familiarity with fewer than two thirds of the faces in a post-experiment questionnaire; and one due to poor compliance with the task instructions leading to lower than chance performance in the final memory test. All participants had normal or corrected to normal vision and no color-blindness. They were screened to ensure they had no neurological conditions or any other contraindications for MRI. Participants were paid 280 shekels (equivalent to ∼$77) for the study. They were recruited from the Hebrew University of Jerusalem community and provided written informed consent prior to participating in the experiment, in a manner approved by the Tel Aviv Sorasky Medical Center Ethics Committee and The Hebrew University institutional review board.

### Materials

Twelve faces of famous women and thirty-six faces of novel women were used in this study. We used famous faces as our manipulation of prior knowledge because famous faces were shown to elicit a rich representation of previous knowledge (for reviews, see Gobbini & Haxby, 2007; Natu & O’Toole, 2011) and were used in previous studies examining prior knowledge influences on new associative learning (Bein et al., 2019; Liu et al., 2016). We further followed a previous multivariate fMRI study using female faces to capture the representation of knowledge about faces (Verosky, Todorov, & Turk-Browne, 2013). The famous faces depicted well-known international and Israeli individuals from a range of fields, including politicians, musicians, actors, and fashion models. An extensive pilot study verified that these faces were indeed familiar to the Hebrew University population and that the participants could identify them by name and provide details about them. The novel faces were obtained from the Web and included foreign corporate executives, actors, and models that were unfamiliar to our Israeli participants, while controlling for factors such as attractiveness and image quality. Of the 36 novel faces, 12 were selected to match the famous faces with respect to age. For convenience, we refer to the famous faces and the 12 matched novel faces as *A-faces*. The remaining 24 novel faces are referred to as *B-faces*. The B-faces were paired with the famous and novel A-faces. The pairing of each of the 24 novel B-faces with each of the 12 famous or 12 novel A-faces was random for each participant, although the pairings were fixed within the experiment (i.e., the same pair appeared in all the repetitions). To enable the associative learning task (see Procedure section below), we added 6 female faces (3 famous) and 12 novel male faces that comprised mixed-gender pairs.

All the stimuli were color photos of faces presented in the center of a grey rectangle that was 290 pixels (width) by 320 pixels (height). The screen resolution was set to 1024*768. To further control for potential visual differences between the pictures, we equated pixel-wise similarity (the correlation across the pixel values between the stimuli; Peelen & Caramazza, 2012; Thierry, Martin, Downing, & Pegna, 2007). Since we used color images, we correlated the RGB values of the stimuli with one another, each color layer separately, and averaged the correlation coefficient of each pair of stimuli across the three layers. We also computed pixel-wise similarity using greyscale versions of the images, in accordance with previous studies. Overall, the correlation values did not differ between the two measures, indicating the viability of pixel-wise color similarity as a measure of pixel-wise similarity (Bein et al., 2019).

A few types of pixel-wise similarity were equated across the stimuli. First, we ascertained that the famous faces were equally distinct from one another and from their matched novel A-faces. We computed the pixel-wise similarity between each of the famous faces and the remaining famous faces and followed the same process for the novel A-faces. We obtained similar means and standard deviations for pixel-wise similarity across the conditions. Next, we verified that on average, the visual similarity between the famous A-faces and B-faces was equal to that of the novel A-faces and B-faces. To that end, we computed the pixel-wise similarity of all the famous faces with all the novel B-faces and the similarity of the novel A-faces with all the novel B-faces. Once again, we obtained similar means and standard deviations for pixel-wise similarity across the conditions.

### Procedure

The experiment started with pre-learning scans, enabling us to capture the multivariate activity pattern of each face alone prior to learning. This was followed by an associative learning session and post-learning scans, to capture the stimuli patterns after learning. Then, a surprise associative memory test was administered. Critically, testing memory after the post-learning scans allowed us to measure post-learning representations without interference from probing of memory. The test was followed by an irrelevant task that was not analyzed. All phases were performed in the scanner and each phase was preceded by detailed instructions and a few practice trials. Upon completion of all tasks, the participants left the scanner and completed a knowledge questionnaire about the faces that appeared in the experiment and a short debriefing session.

#### Pre/post-learning scans

The pre- and post-learning scans were identical (Kim et al., 2017; Schapiro et al., 2012; Schlichting et al., 2015). All faces that appeared in the learning phase appeared in these scans. In each trial, a face appeared alone at the center of the screen for 1 s. Trials were jittered with .5-7.5 s of a fixation-cross baseline, with an interval of .5 s, using optseq2 (https://surfer.nmr.mgh.harvard.edu/optseq/; Dale, 1999). Participants were asked to indicate by pressing a button whether the person appearing on the screen was male or female.

Each phase (pre and post) was divided into two scans; in each scan, each face appeared three times. The order of stimulus presentation was pseudo-randomized to maintain low autocorrelations between regressors and to ensure that two faces that appeared as a pair in the associative learning task appeared with a minimal gap of two stimuli in the pre/post scans, to prevent additional learning during these scans. To create the pseudo-randomized order, placeholders of stimuli were fixed, e.g., a certain face appeared in locations 20, 150, and 180, comprising the regressor of that face. Regressors were then paired such that two faces that would be associated later would each appear in one of the regressors of the pair (e.g., placeholders 20, 150, and 180 were paired with placeholders 40, 105, and 240). We had two of these fixed orders (determined by simulations to ensure low correlations), one for each scan, and the order of the scans was counterbalanced across participants. To counterbalance the conditions, the pairs of placeholders within each scan were divided into two groups of 12 place-holder pairs. The allocation of famous and novel A-faces (and the corresponding B-faces) to placeholder groups was counterbalanced across participants. Within each placeholder group, the allocation of placeholders to either A or B faces rotated across participants. The allocation of the stimuli to placeholders was randomized within each condition (famous/novel and A/B-face) for each participant. Critically, the pre and post scans were identical within each participant (we also repeated the same order of scans) and the pattern similarity before learning was subtracted from the pattern similarity after the learning. Thus, differences in pattern similarity cannot be attributed to differences in the correlations between regressors (Shapiro et al., 2012). All faces appeared once before a new cycle of repetition began. Although we analyzed only female faces that appeared in the same-gender pairs during learning, the stimuli from the mixed-gender pairs were included in the pre/post scans as well, to equate familiarity of the stimuli during the associative learning task, and to enable the male/female gender task during the pre/post scans. The placeholders of these additional faces were fixed across participants to distribute the males throughout the task, but the allocation of the faces to placeholders was randomized for each participant.

#### Associative learning task

Participants were presented with pairs of faces that were composed of either a famous and a novel face (Prior Knowledge, PK) or two novel faces (No Prior Knowledge, n-PK). In each trial, the faces were presented at the center of the screen. The faces were presented sequentially and not simultaneously to prevent participants from fixating on one face more than the other. Each trial included a double repetition of the pair (A-B-A-B), with each face appearing on the screen for 500 ms^5^ and an inter-stimulus interval (ISI) of 100 ms. A fixation cross appeared at the end of each trial for 600 ms. As before, trials were further jittered with .5-7.5 s fixation-cross baseline, with an interval of .5 s (Dale, 1999). The participants had to indicate by pressing a button whether the two faces were two females or a male and a female and were instructed to respond as quickly and as accurately as they could.

Throughout the learning phase, each pair was repeated 12 times (12 repetitions of A-B-A-B trials). This task was divided into four scans, each of which included three presentations of all pairs. Each cycle of repetition included all 24 experimental pairs and the additional 6 different-gender pairs, which also repeated 12 times throughout the experiment. In these filler trials, the male always appeared second (as a B-face). To allow enough males for the pre/post-learning scans, each female was paired with two males, and these appeared alternatively (such that each male appeared six times in total during the associative learning task). In each cycle, the order of stimuli was pseudo-randomized in a similar manner to the pre/post-learning scans: placeholders were fixed and divided into two groups (for the two conditions, PK and n-PK). The allocation of the groups was counterbalanced across participants. Within each group, the allocation of a specific pair to the placeholders was randomized for each participant. We had four such fixed orders (determined again by simulations to ensure low correlations between regressors), one for each scan, and the order of the scans was randomized for each participant. The placeholders of the female-male pairs were fixed to allow distribution of these trials throughout the task. All pairs appeared once before a new cycle of repetition began.

#### Associative memory tes

Upon completing the associative learning phase, participants performed the post-learning scans. Then, a surprise memory test was given. In each trial, participants were presented with an A-face that appeared at the top of the screen (either famous or novel) and three B-faces that appeared at the bottom of the screen, one of which had been paired with the A-face during the learning session. All three faces were intra-list within the pair type (i.e., if an A-face was famous, the two distractors were B-faces that appeared with other famous faces). The allocation of the distractors was pseudo-randomized such that a triad of the same B-faces could not appear twice throughout the test. One third of the B-faces appeared as targets in their first presentation, one third appeared as targets in their second presentation, and one third appeared as targets in their third presentation. Within each condition (PK/n-PK), the location of the target was equally divided between the three possible locations, and each B-face appeared once as a target and twice as a distractor.

In each trial, participants were asked to choose the B-face that had appeared with the A-face during the learning session. After a face was chosen, the other two faces disappeared, and participants were asked to make a three-level confidence judgment (sure, probably, or maybe, corresponding to high, medium, or low confidence that the faces appeared together, respectively). Both stages of each trial were self-paced, but each stage was limited to 10 s. A 500 ms fixation cross appeared between trials. The order of the trials was randomized such that no more than two trials of either the PK or the n-PK condition appeared consecutively. Within each condition, trials were randomized and B-faces were allocated as distractors such that no face would appear in two consecutive trials, either as target or as distractor.

#### Knowledge questionnaire

After scanning, participants completed a knowledge questionnaire. All faces appeared one after the other, and subjects had to say whether they knew their names or were familiar with them before the experiment. They were additionally asked to rate how many facts they knew about the person whose face was presented. Since we piloted the famous faces to ensure that people had knowledge about them, this questionnaire was only meant to crudely assess the knowledge of the specific participant. Thus, we excluded subjects that did not recognize (i.e., could not provide the person’s name or reported that the person was not familiar to them) over a third of the famous people in the study (two participants). We further excluded from all analyses specific famous faces that were not familiar to a particular participant (six participants each had one face excluded, a different famous face across these participants). Then, participants were debriefed and asked whether they had suspected that there would be a memory test or tried to memorize the pairs during the learning phase.

### fMRI parameters and preprocessing

Participants were scanned in a 3T Simens Prisma scanner. The experiment included an MPRAGE anatomical scan (1X1X1mm resolution), a fieldmap scan, and 12 whole-brain T2*-weighted EPI scans (TR=2000 ms, 200-mm*180mm FOV, 64×58 matrix, TE=28, flip angle=77, phase encoding direction: *anterior*-posterior). In each volume, 39 slices were acquired tilted minus 20 degrees of the AC-PC, 3.125*3.125*3.1-mm (width*length*thickness) voxel size, no gap, in a top-down interleaved order. In each of the four sessions of the pre-post task, 366 images were acquired. Each of the four sessions of the learning task included 179 images.

The imaging data were preprocessed using SPM8 (Wellcome Department of Cognitive Neurology) for MATLAB (Mathworks, Natick, MA), FSL (http://www.fmrib.ox.ac.uk/fsl), and in house scripts for the similarity analysis. Images were corrected for differences in slice acquisition timing and realigned to the mean image across all scans to correct for movement. Neither smoothing nor registration to standard space was performed, as all analyses were made in subject space. For group level-analyses of functional connectivity during the learning task, subject-level t-stats maps were smoothed and registered to MNI space (see below).

### Regions of Interest (ROIs)

The hippocampus was defined anatomically for each participant using FSL’s automatic subcortical segmentation protocol (FIRST). The hippocampus was segmented along its long axis by dividing the number of coronal slices in each hemisphere into three sections. The anterior third of the coronal slices was designated as anterior hippocampus, and the posterior third of the coronal slices was designated as posterior hippocampus (Tompary & Davachi, 2017). We further divided the hippocampus a-priori to left and right hemisphere. We examined these four hippocampal ROIs (left/right by anterior/posterior), as previous findings on prior knowledge in the hippocampus do not coincide with respect to the specific locus of influence (Brod et al., 2016; Reggev et al., 2016).The left inferior frontal gyrus (left IFG) and the angular gyrus were defined functionally, based on the group level contrast of PK > n-PK in the PPI analysis detailed below. Then, for representational similarity analyses, we brought the peak voxel to each participant’s native space and constructed a 12-mm sphere (10-mm sphere yielded similar results).

### Representational Similarity Analysis

For each subject, one GLM was constructed for the two pre-learning scans and one for the two post-learning scans. To model the response for each face in each session, the canonical hemodynamic response was convolved with the onset of the three presentations of the face in a session (time derivative regressors were added, as well as a constant for each scan and a 128 s high pass filter). This yielded a beta-value for each stimulus in each of the four scans. We then converted these beta values into t-stats and averaged, for each stimulus, the two t-stats of the pre-learning scans, to obtain the multivoxel activity pattern before learning. The same was done for the two t-stats from the post-learning scans, to obtain the pattern of that face after learning. These values were then Fisher transformed for statistical analysis.

We conducted two types of representational similarity analyses:

1. <u>Memory-related pre-to-post similarity differences:</u> We examined similarity differences that mediated explicit memory by computing for each participant the average similarity difference between pre- and post-learning in each pair type (PK/n-PK), and within each memory outcome. That is, within each pair type, we averaged similarity differences for pairs that were remembered with high confidence in the subsequent memory test (high-confidence included “certain” and “probably” responses; excluding low confidence trials is a common practice used in fMRI studies to exclude guesses, Wagner et al., 1998; One participant with no high confidence hits in the PK condition was removed from this analysis). We then compared this average to the average similarity for pairs that were forgotten (misses), within each pair type. To that end, differences in similarity from pre-to post-were entered into a repeated-measures ANOVA of Pair Type (PK/n-PK) by Memory (Remembered/Forgotten). This ANOVA was followed up by two-tailed paired-sample t-test, addressing simple effects.
2. <u>Asymmetry in representational changes:</u> To assess whether cortical learning was asymmetric (see Introduction), for each pair that appeared together during learning, we subtracted the similarity of the A-face post-learning to that of the B-face pre-learning, from the similarity of the B-face post-learning to the A-face pre-learning (Schapiro et al., 2012). This gave us a measure of how much more similar the B-face representation after learning became to that of the A-face before learning, as compared to the extent to which the A-face became similar to the B-face. We then computed the average for each participant across all pairs per pair type (PK/n-PK). As control baseline, we further computed the same asymmetry index for shuffled pairs. Shuffled pairs were obtained by pairing each A-face with all other B-faces that did not appear with that A-face during learning, but critically, appeared in the same pair type. We then averaged for each participant all shuffled-pairs per pair type to obtain a baseline asymmetry index. Asymmetry was compared to 0 or to shuffled pairs using two-tailed paired-sample t-tests.

### Functional connectivity analysis

We conducted PPI analysis (SPM8 gPPI toolbox, McLaren et al., 2012) during the associative learning task, with the left anterior hippocampus ROI as a seed region (in pre-to post-learning similarity differences, this region showed separation for remembered PK pairs, but similarity for remembered n-PK pairs, see Results). Thus, for each participant, the times-series of the left anterior hippocampus during the associative learning task was used as the physiological regressor. To reflect the nature of our task, the psychological regressors included a regressor for all pairs in each pair type, in each cycle of repetition (a total of 24 regressors, 12 repetitions by PK/n-PK pair type). The psychophysiological regressors were the interaction of each psychological regressor with the physiological regressor. As before, a high-pass filter of 128 s and constant scan regressors were added for each scan. We then computed, for each participant, the contrast of all repetitions of the PK pair type versus all repetitions of the n-PK pair type. Analyses were performed in each participant’s native functional space. The resulting t-maps were then smoothed (8-mm HFWM kernel) and registered to the MNI space for group-level analysis. At the group level, the PK versus n-PK contrast was compared to zero using a one-sample two-tailed *t*-test. The resulting t-map was thresholded at a voxel-level of *p* < .005 due to low power in PPI designs (O’Reilly, Woolrich, Behrens, Smith, & Johansen-Berg, 2012), accounting for the reduced voxel-level threshold by maintaining a cluster level threshold of *p* < .05 (Brod et al., 2016; resulting in cluster size > 61 voxels, Monte Carlo simulations, Slotnick, Moo, Segal, & Hart, 2003).

## Footnotes

1 Note that this logic is valid only in one direction of inference. Distinct representations, if observed, will provide confirmatory evidence for the interference resolution hypothesis. Naturally, finding such separation cannot guarantee that distinct representations mediate reductions in interference.

2 The interaction survived Bonferroni correction for 4 ROIs (p < .0125): left/right hemisphere by anterior/posterior hippocampus. Main effect of Prior Knowledge in the left anterior hippocampus: *F*_(1,17)_ = 4.09, *p* = .059. Interactions or main effects in other hippocampal ROIs: all *F*’s _(1,17)_ < .88, *p*’s > .36, but the main effect of Prior Knowledge in the right posterior ROI: *F*_(1,17)_ = 2.82, *p* = .11.

3 No medial-prefrontal cortex (mPFC) clusters emerged in this analysis. While not all prior knowledge studies report medial-prefrontal findings (for a review, see Gilboa & Marlatte, 2017), a recent study did find slightly higher hippocampus-mPFC connectivity when participants associated pairs of famous faces and houses, as compared to novel faces and houses (Liu et al., 2016). Given the broad interest in hippocampus-mPFC interactions (Eichenbaum, 2017; Preston & Eichenbaum, 2013), and specifically in relation to prior knowledge (Bein et al., 2014; Gilboa & Marlatte, 2017; van Kesteren, Fernandez, et al., 2010; van Kesteren et al., 2012), we examined whether a mPFC cluster would emerge using a more liberal threshold of *p* < .01, voxel-level. Indeed, a region showing higher connectivity in PK pairs compared to n-PK pairs emerged in the ventral and anterior part of the mPFC ([2, 52, −24], 163 voxels). The opposite contrast did not reveal any mPFC cluster at this statistical threshold.

4 We also examined the angular gyrus (AG), which was recently proposed to be a hub, binding aspects of schematic knowledge and mediating schema influences on encoding (Bein, Reggev, & Tompary, 2018; Gilboa & Marlatte, 2017; van der Linden, Berkers, Morris, & Fernández, 2017; I. C. Wagner et al., 2015). We found no initial similarity differences, with or without respect to subsequent memory (ANOVA’s of Prior Knowledge by Paired/Shuffled, or of Prior Knowledge by Memory, *F*’s < 2.08, *p*’s > .16). We thus did not proceed to investigate whether similarity differences were asymmetric, as we had no evidence that they had occurred.

5 This presentation time ensured recognition of the famous faces (Bentin & Deouell, 2000; Neumann & Schweinberger, 2008; Tacikowski, Jednoróg, Marchewka, & Nowicka, 2011).

