## Supplementary Information for "The direction of associations: prior knowledge promotes hippocampal separation, but cortical assimilation"

*Behavioral results*

During the associative learning task, participants viewed pairs of faces. Some pairs contained a famous and a novel face (prior knowledge, PK) and some pairs included two novel faces (no prior knowledge, n-PK). The pairs were repeated 12 times in cycles, and each cycle included all pairs in random order. For each pair, participants judged whether both faces were of the same gender (experimental trials were always of the same gender; we interspersed mixed-gender pairs as fillers to make the task possible, see Methods). Participants performed at ceiling for the gender judgements across all pair types and repetition cycles (PK and n-PK, same- and mixed-gender pairs; all > 96%). The mixed-gender pairs were not further analyzed. We further excluded from all analyses PK pairs in which participants did not recognize the famous face in the post-experiment knowledge questionnaire (for six participants, one pair was removed; the pairs included different famous faces across these participants). For analysis of the reaction times (RTs) during learning, we first excluded all incorrect responses and trials in which participants responded more than once. We further excluded, for each participant, trials in which RTs deviated more than 3 standard deviations from the mean in each cycle of repetition. On average, 8.42 and 5.53 responses (6% and 4%) were excluded for PK and n-PK pairs, respectively. Participants demonstrated learning, as indicated by faster RTs towards the end of learning than early in learning (Figure S1).

For statistical analysis, single-trial RTs were entered as the predicted variable to general linear mixed models (gLMM, as implemented by the glmer function, lme4 package in R; Bates, Mächler, Bolker, & Walker, 2014) with inverse Gaussian distribution as the linking distribution function (Lo & Andrews, 2015). Model comparisons were used to examine the effects of Pair Type (PK/n-PK), Repetition (1-12 as a continuous variable), and their interaction. All models included a random intercept per participant. We found that Repetition significantly predicted the decrease in RT, indicating learning. Specifically, a model including the Repetition and Pair Type factors significantly outperformed a model that included only Pair Type, indicating that Repetition significantly explained variance in RTs (*χ*^2^ = 105.58, *p* < .0001, AIC difference: 104, BIC difference: 97). Repetition significantly explained RTs when a simpler model was conducted, which included Repetition as a single fixed-effect, and when this model was compared to a null-model that only included a random intercept per participant and no fixed effect (*χ*^2^ = 105.3, *p* < .0001, AIC difference: 103, BIC difference: 97). There was also a small effect of Pair Type, suggesting that participants responded to PK pairs slightly faster than to n-PK pairs. This effect was significant when comparing the full model, including Pair Type and Repetition, versus a model including only Repetition (*χ*^2^ = 3.86 , *p* < .05, with a minor AIC difference of 2, but BIC difference of -5), and marginally significant when comparing the simpler model, including only Pair Type, to the null model (*χ*^2^ = 3.58 , *p* = .058, with a minor AIC difference of 1, but BIC difference of -5). There was no Pair Type by Repetition interaction, as indicated by comparing a model with Pair Type, Repetition, and their interaction, to the same model without the interaction term (*χ*^2^ = .18, *p* > .67).

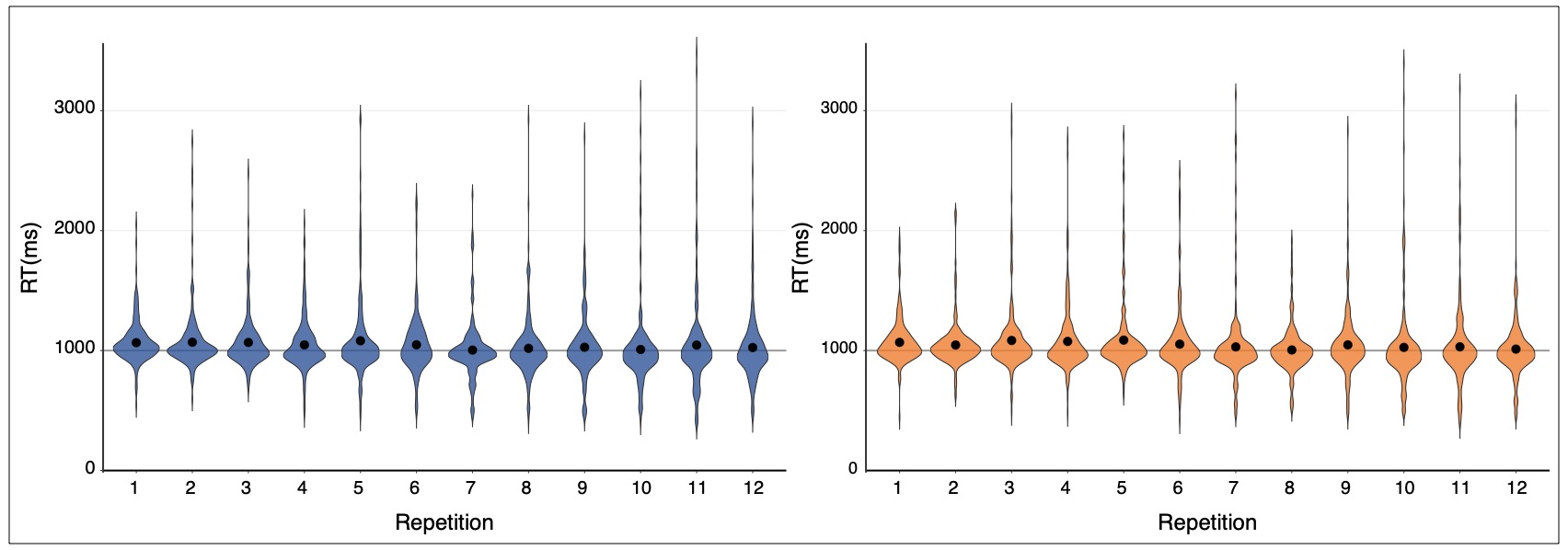
During the pre- and post-learning scans, participants made male/female judgments for single faces that appeared in the associative learning task. We term faces that appeared as the first face in a pair during the learning task A-faces and faces that appeared second B-faces. Accuracy was at ceiling during the pre-scan and the post-scan. On average, participants responded to all faces with 96-99% accuracy during both pre-learning and post-learning scans (rates for all types of faces, e.g., famous faces and novel faces did not differ between pre- and post-learning scans.)

*Figure S1.* Reaction times (RT) during the associative learning task in each repetition cycle. Left, in blue: RT for the prior knowledge pair type (PK). Right, in orange: RT for the no prior knowledge pair type (n-PK). The black dots indicate mean RT.

*Imaging Results*

*Functional connectivity with the left anterior hippocampus*

|  | *MNI coordinates* | | |  |  |
| --- | --- | --- | --- | --- | --- |
| *Region* | *x* | *y* | *z* | *Z value* | *Num. Voxels* |
| Dorsomedial prefrontal cortex | 6 | 36 | 50 | 3.92 | 1042 |
| L orbitofrontal cortex | -24 | 48 | -14 | 3.67 | 170 |
| L middle temporal gyrus | -44 | 0 | -30 | 3.66 | 568 |
| R superior frontal gyrus | 10 | 62 | 28 | 3.50 | 62 |
| R inferior frontal gyrus | 42 | 32 | -22 | 3.38 | 269 |
| L inferior frontal gyrus (ventral) | -48 | 30 | -16 | 3.27 | 87 |
| L inferior frontal gyrus (dorsal) | -52 | 30 | 10 | 3.00 | 70 |
| L angular gyrus | -42 | -48 | 26 | 3.19 | 126 |
| Cerebellum | -24 | -88 | -42 | 2.94 | 67 |

*Supplementary Table 1.* Regions demonstrating greater functional connectivity (gPPI) with the left anterior hippocampus for prior knowledge (PK) pairs compared to no prior knowledge (n-PK) pairs during associative learning. No region was observed in the opposite contrast (see main text, Methods, Results, and Figure 3).

*Learning-related similarity differences in the left inferior frontal gyrus*

We targeted asymmetric changes in the representational patterns in the left inferior frontal gyrus (IFG) for famous and novel faces (see main text). However, asymmetry in representational changes from before to after learning only makes sense if some representational change indeed happened from before to after learning. To that end, we first asked whether there was evidence for learning-related similarity changes from pre- to post-learning in the left IFG between. For each pair type (PK/n-PK), we compared pairs of items that appeared together during the learning task to the shuffled pairs baseline. The shuffled pairs baseline was obtained by pairing each A-face with all other B-faces of the same pair type. Pre and post similarity differences were computed for each such pairing and averaged across all pairs, like in the associated pairs. Note that this within-condition shuffling controlled for any differences the B-faces might have had due to their appearance with famous versus novel pairs, since for each pair type, all faces in the shuffled baseline appeared with A-faces of the same pair type. Similarity differences for PK and n-PK pairs were thus computed and compared to similarity for shuffled pairs. This comparison allowed us to specifically examine changes due to associative learning, controlling for similarity differences due to item familiarity or type of pair (Schapiro et al. 2012). Similarity differences in the left IFG were then Fisher-transformed and entered into a repeated-measures ANOVA with Prior Knowledge (PK/n-PK) and Pairing (paired/shuffled), revealing a main effect of Pairing (*F*_(1,18)_ = 11.44, *p* = .003, η*_p_*^2^ = .32), and no main effect of Prior Knowledge or Prior Knowledge by Pairing interaction (*F*’s < 1.2, *p*’s > .29). Pairwise comparisons showed a highly significant increase in similarity in PK pairs (PK, paired vs. shuffled: *t*_(18)_ = 4.1, *p* = .0007, Cohen’s *d* = .94). N-PK paired faces also became more similar compared to shuffled baseline, but not significantly so (*t*_(18)_ = 1.16, *p* = .26). While the similarity differences were specific to paired faces compared to shuffled faces and thus clearly indicated learning, they were not related to subsequent memory in the current study, as no main effect or interaction was obtained in the Prior Knowledge (PK/n-PK) by Memory (remembered/forgotten) ANOVA (all *F*’s < 2.46, *p*’s > .13).

*Control for univariate activation*

To rule out the possibility that univariate activation accounted for our similarity findings, we included univariate activation during the pre and post similarity scans as a factor in multiple regression models. For each participant in each of the relevant ROIs, the faces’ t-maps were binned and averaged in each pair type (PK/nPK) and by A/B-faces. For the memory analysis, maps were further divided to remembered pairs (high-confidence hits) and forgotten pairs (misses), paralleling the similarity analysis.

*Left anterior hippocampus*

We controlled for univariate activation in the Prior Knowledge by Memory interaction by including univariate activation in models implemented via ANOVA (aov function in R, stats package). Similarity differences per participant in each of the four bins (PK/n-PK by remembered/forgotten) were the explained variables. As explaining variables of interest, we included Prior Knowledge (PK/n-PK), Memory (Remembered/Forgotten), and their interaction. Since our similarity measures are a difference between post and pre-similarity, in the first model we controlled for the difference in univariate activation between post and pre similarity, by including in the model these differences in each of the four bins (PK/n-PK by remembered/forgotten), for both A-faces and B-faces. The models further included a within-participant error term for each of the factors of interest (i.e., participant/Prior Knowledge*Memory). The Prior Knowledge by Memory interaction was significant (*F*_(1,15)_= 9.44, *p* = .008, η*_p_*^2^ = .39). In a second model, instead of activation differences, we included activation for pre and post scans separately (four variables: A/B face by pre-/post-learning). Again, a significant interaction was obtained (*F*_(1,13)_= 10.05, *p* = .007, η*_p_*^2^ = .43). We proceeded to the simple effect obtained between remembered PK pairs and remembered n-PK pairs. We repeated the same models as before, controlling for the difference in univariate activity between pre- and post-learning in one model, and each phase separately in another model, but including only values of remembered pairs (for similarity and univariate data). In both models, an effect of Prior Knowledge was revealed (control for pre-post differences: *F*_(1,15)_= 9.90, *p* = .007, η*_p_*^2^ = .40; each phase: *F*_(1,13)_ = 9.10, *p* = .01, η*_p_*^2^ = .41). Then, to test for the simple effects of Memory within each pair type, we repeated the same two models as before, but now taking first only PK pairs, then only n-PK pairs (remembered vs. forgotten pairs, now only a Memory variable was included in addition to the univariate variables). The effect of Memory was significant in all four models (significant in PK pairs, pre-post differences: *F*_(1,15)_ = 6.04, *p* = .03, η*_p_*^2^ = .29; each phase: *F*_(1,13)_ = 5.98, *p* = .03, η*_p_*^2^ = .32; marginally significant in n-PK pairs: *F*_(1,15)_ = 3.38, *p* < .09, η*_p_*^2^ = .18; each phase: *F*_(1,13)_ = 3.64, *p* < .08, η*_p_*^2^ = .22).

*Left inferior frontal gyrus*

The asymmetry measure in PK pairs was different from 0, as well as from the asymmetry in shuffled pairs. Thus, in the current analyses, we wished to control for univariate activation for these two comparisons. As before, we accounted for univariate activation of A- and B-faces. Note, however, that both paired and shuffled pairs were computed using the same A and B faces, but with these faces paired differently to compute the asymmetry. Therefore, there was no separate univariate activation for A- and B-faces in paired and shuffled pairs, as both included the same faces. Thus, to allow us to examine whether the difference in asymmetry between paired and shuffled faces would holds when controlling for univariate activation of the A and B faces, we needed one asymmetry measure per participant. To that end, for each participant, we subtracted the asymmetry measure in the shuffled faces baseline from the asymmetry measure for paired faces. This difference measure was then taken as the explained variable when comparing the difference in asymmetry for paired versus shuffled pairs. In additional models, we also examined the raw asymmetry measure (to control for univariate activity for the comparison against 0). As the explaining variables, we included as before either the pre-post differences for A and B faces, or each of the pre and post phases separately. This yielded at total of four models (difference from 0/difference from shuffled pairs by activation difference/each phase separately). The linear regression models were evaluated using the lm function in R (stats package; note that the explained variable is already a within-participant difference measure, hence there is no need to include a within-participant error term, as was done above). In all four models, the asymmetry measure was significant (the group intercept, paired-shuffled, pre-post differences: *β* = 0.029, *t*_(16)_ =3.66, *p* = .002; each phase: *β* = 0.037, *t*_(14)_ = 4.46, *p* < .001; paired-0, pre-post differences: *β* = 0.026, *t*_(16)_ =2.65, *p* = .017; each phase: *β* = 0.029, *t*_(14)_ = 3.23, *p* = .006).

Taken together, these analyses confirm that our similarity findings are unlikely to be attributed to differences in univariate activation.
